# NETMAGE: a humaN-disEase phenoType MAp GEnerator for the Visualization of PheWAS

**DOI:** 10.1101/2020.10.27.357103

**Authors:** Vivek Sriram, Manu Shivakumar, Sang-Hyuk Jung, Lisa Bang, Anurag Verma, Seunggeun Lee, Eun Kyung Choe, Dokyoon Kim

**Affiliations:** Genomics and Computational Biology Graduate Group, Perelman School of Medicine, University of Pennsylvania, Philadelphia, PA 19104, USA; Department of Biostatistics, Epidemiology & Informatics, Perelman School of Medicine, University of Pennsylvania, Philadelphia, PA 19104, USA; Samsung Advanced Institute for Health Sciences and Technology (SAIHST), Sungkyunkwan University, Samsung Medical Center, Seoul, Republic of Korea; Ultragenyx Pharmaceutical, Novato, CA 94949, USA; Department of Genetics, Perelman School of Medicine, University of Pennsylvania, Philadelphia, PA 19104, USA; Graduate School of Data Science, Seoul National University, Seoul, South Korea; Department of Surgery, Seoul National University Hospital Healthcare System Gangnam Center, Seoul, 06236, South Korea; Institute for Biomedical Informatics, University of Pennsylvania, Philadelphia, PA 19104, USA

## Abstract

**Summary:** Given genetic associations from a PheWAS, a disease-disease network can be constructed where nodes represent phenotypes and edges represent shared genetic associations between phenotypes. To improve the accessibility of the visualization of shared genetic components across phenotypes, we developed the humaN-disEase phenoType MAp GEnerator (NETMAGE), a web-based tool that produces interactive phenotype network visualizations from summarized PheWAS results. Users can search the map by a variety of attributes, and they can select nodes to view information such as related phenotypes, associated SNPs, and other network statistics. As a test case, we constructed a network using UK BioBank PheWAS summary data. By examining the associations between phenotypes in our map, we can potentially identify novel instances of pleiotropy, where loci influence multiple phenotypic traits. Thus, our tool provides researchers with a means to identify prospective genetic targets for drug design, contributing to the exploration of personalized medicine.

**Availability and implementation:** Our service runs at https://hdpm.biomedinfolab.com. Source code can be downloaded at https://github.com/dokyoonkimlab/netmage.

**Supplementary information:** Supplementary data and user guide are available at *Bioinformatics* online.

## 1. Introduction

Phenome-Wide Association Studies (PheWASs) identify genetic associations across numerous phenotypes through the combined analysis of genomic and phenotypic data. Through the analysis of PheWAS results, we can identify instances of novel pleiotropy among diseases, where a single gene affects multiple unrelated phenotypic traits^1^. Visualization of the associations between genes and phenotypes can provide clinicians with an effective way to understand connections across diseases. A variety of tools currently exist for the representation of PheWAS results, including “PheWAS-Me,”^2^ “PleioNet,”^3^ “ShinyGPA,”^4^ “PheGWAS,”^5^ “Multi PheWAS Viewer,”^6^ and “PheWeb”^7^. However, such packages are unable to specifically present interactive, searchable network displays of genetic associations between phenotypes given any sort of input summary PheWAS data (Supplementary Data). The creation of a “disease-disease network” (DDN), where phenotypes are represented by nodes and shared genetic associations are represented by edges, would help researchers and clinicians gain a deeper understanding of the underlying genetic architecture across diseases^8^. Thus, we developed the humaN-disEase phenoType MAp GEnerator (NETMAGE), a web-based tool that produces interactive phenotype networks from input summary PheWAS data. NETMAGE allows researchers to visualize and interact with the genetic connections between phenotypes for any dataset.

## 2. Implementation

We used InteractiveVis^9^, an interactive network framework built over sigma.js, as a base for the implementation of NETMAGE. We developed a web interface for the generation of network visualizations. A primary consideration in NETMAGE was allowing for the definition of additional attributes for search and filtration in the network. Thus, we extended the framework to automatically parse any additional attributes provided by the user in the input file and turn them into options for filtration and search. Furthermore, the generated interactive network can be viewed online or downloaded. The downloaded files can be used to host the interactive network on any other host.

## 3. Platform Description

The DDNs generated from NETMAGE come with a variety of features that allow users to explore genetic associations between phenotypes:

1. Uploading PheWAS Results – we provide a script to generate edge and node map files from input PheWAS results and instructions for how to convert these files into a network JSON file through Gephi. NETMAGE will then process this Gephi output file to generate a corresponding visualization.
2. Node Selection – clicking on a node will highlight the node and all of its first-degree neighbors. A variety of default attributes will be presented on the right side of the webpage (Fig. 1). Other custom attributes can also be defined by the user.
3. Search – users can search the map for relevant phenotypes based upon any attribute defined, such as phenotype name, phenotype ID, SNP name, node degree, and other parameters.
4. Highlighting – groups of nodes within the same disease category can be highlighted to visualize associations within groups. These categories are established solely by the input dataset.

**Figure 1.**
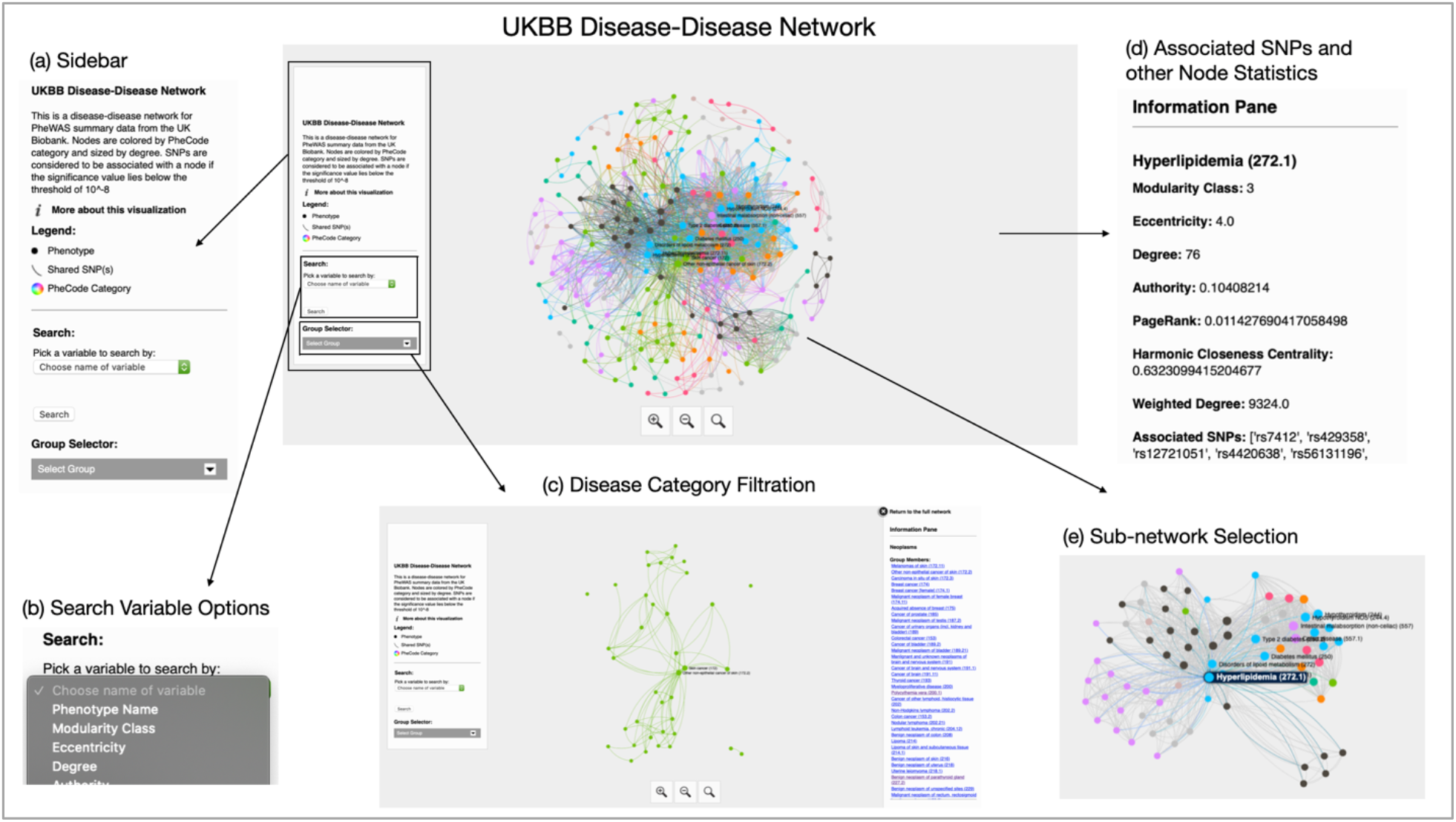
A depiction of the NETMAGE Visualization tool. (a) The sidebar of the visualization gives detail on the map. It also includes a search dropdown as well as a group selector dropdown menu. (b) Variables are automatically read from the input data and included as options for search. (c) All nodes within a single disease category can be visualized at once. All phenotype names in the disease category are also displayed to the right of the visualization. (d) Associated SNPs, connected phenotypes, and a variety of network statistics are presented to the right of the window when a node is selected. (e) Selection of a node also reduces the nodes displayed in the map to only the chosen node and its direct connections.

The user guide can be found in Supplementary Data.

## 4. Case Study – UKBB

As a demonstration of the capabilities of NETMAGE, we applied our software to SAIGE-analyzed UK Biobank (UKBB) PheWAS data^10^. These data included information on 400,000 Caucasian British samples, 1403 phenotypes expressed in terms of PheCodes, and 28 million variants. The “SAIGE” R package was applied to these data to produce p-values of association between SNPs and diseases^11^.

To ensure a manageable number of nodes and edges in our resulting map, these data were filtered using three criteria: 1) p-value threshold of 1×10^-8^, 2) minor allele frequency of 0.01, and 3) case count of 200. Using the filtered data, an adjacency matrix was calculated using the connections between diseases, constituting the “edge map.” Associated SNPs for each phenotype were added as an attribute of the disease nodes to PheWAS Catalog^12^ data to make the “node map” of the data. The software application Gephi^13^ was used to construct a network from these maps, establishing coordinates to a two-dimensional space for our phenotypes. The details about node attributes and disease categories are described in Supplementary Data. Our final map included 260 unique phenotypes and 2,642 genetic associations. 678 (25.66%) of the edges occurred within PheCode categories, while 1964 (74.34%) edges occurred across them. The map is hosted at https://hdpm.biomedinfolab.com/ukbb/.

Our map correctly displays previously identified instances of potential pleiotropy from the UKBB, including rs544873’s association with pulmonary heart disease, phlebitis and thrombophlebitis, hemorrhoids, circulatory disease, and diverticulosis, and rs925488’s association with thyroid cancer, nontoxic nodular and multinodular goiter, and hypothyroidism^14^. Our map also appropriately shows associations for the SNP rs11066309 with hypertension, ischemic heart disease, myocardial infarction, coronary atherosclerosis, and hypothyroidism^15^, and associations for the SNP rs780094 with diabetes and lipid metabolism.^16^

## 5. Discussion

### 5.1 Strengths of NETMAGE

Key strengths of NETMAGE include the automated creation of maps from user input for the visualization of a multitude of datasets, searchability of network maps by SNP for the search for potential pleiotropy, and interactivity with network nodes, giving users the opportunity to focus on specific genetic associations by visualizing subsets of the map. Our map also offers the ability to peruse edges for the identification of pleiotropy.

### 5.2 Conclusions and Future Work

NETMAGE is a web-based toolkit for the interactive visualization of PheWAS summary data, improving the ease of visualization of genetic associations across diseases. In terms of future directions for our software, we would like to introduce functionality to make “gene-gene networks” that depict associations between genes based upon shared effects on phenotypes. We would also like to incorporate methods to compare networks to one another. By running our software on additional datasets such as the Penn Medicine Biobank and comparing resulting output, we could evaluate which genetic associations may be false positives, and which ones are true novel instances of pleiotropy. Ultimately, NETMAGE will give clinicians insight into the underlying genetic architecture of diseases. We hope that this software will contribute to new potential discoveries in personalized medicine and drug development.

## Funding

This work has been supported by the NIGMS R01 GM138597.

## Supplementary Data

### 1. Comparison of PheWAS Visualization Toolkits

| **Software Name** | **Allows users to upload desired PheWAS results for analysis** | **Allows for interactive investigation of pleiotropy** | **Generates a network visualization of genetic associations between phenotypes** | **Allows users to search and create subsets of any produced networks by disease, by SNP, or by other network statistics** |
| --- | --- | --- | --- | --- |
| PleioNet |  | **x** | **x** | **x** |
| ShinyGPA | **x** | **x** |  | **x** |
| PheGWAS | **x** | **x** |  | **N/A** |
| PheWAS-Me | **x** | **x** |  | **x** |
| Multi PheWAS Viewer |  | **x** | **x** |  |
| PheWeb | **x** | **x** |  | **N/A** |
| NETMAGE | **x** | **x** | **x** | **x** |

### 2. Node Attributes and Disease Categories available for the UKBB Network

Network statistics that can be viewed in the UKBB network include: modularity class, eccentricity, degree, authority, page rank, harmonic closeness centrality, weighted degree, closeness centrality, hub, component ID, eigenvector centrality, clustering coefficient, betweenness centrality, and number of triangles. Node labels are displayed in the initial visualization for phenotypes with the highest degrees.With the UKBB data, options for disease groups include: Digestive, Hematopoietic, Circulatory System, Dermatological, Endocrine/Metabolic, Genitourinary, Sense Organs, Neoplasms, and Musculoskeletal.

### 3. NETMAGE User Guide

#### Availability and implementation

Our service runs at https://hdpm.biomedinfolab.com. Source code can be downloaded at https://github.com/dokyoonkimlab/netmage

#### Creating “data.json” given PheWAS summary data

This first section of the guide provides a description of how to take your input PheWAS data and convert it into ***edge*** and ***node maps***, and then apply ***Gephi*** to generate ***data.json*** from these maps. ***Data.json*** can then be uploaded into NETMAGE to create a corresponding disease-disease network.

#### a) Edge Map

a comma-separated-value file including all established edges within our disease-disease network. This file should include a column for “Sources” and a column for “Targets” respectively, specifying the start and end of each edge in our map. Other columns, such as edge weight or shared SNPs, can optionally also be included in the file.

**Figure 1.**
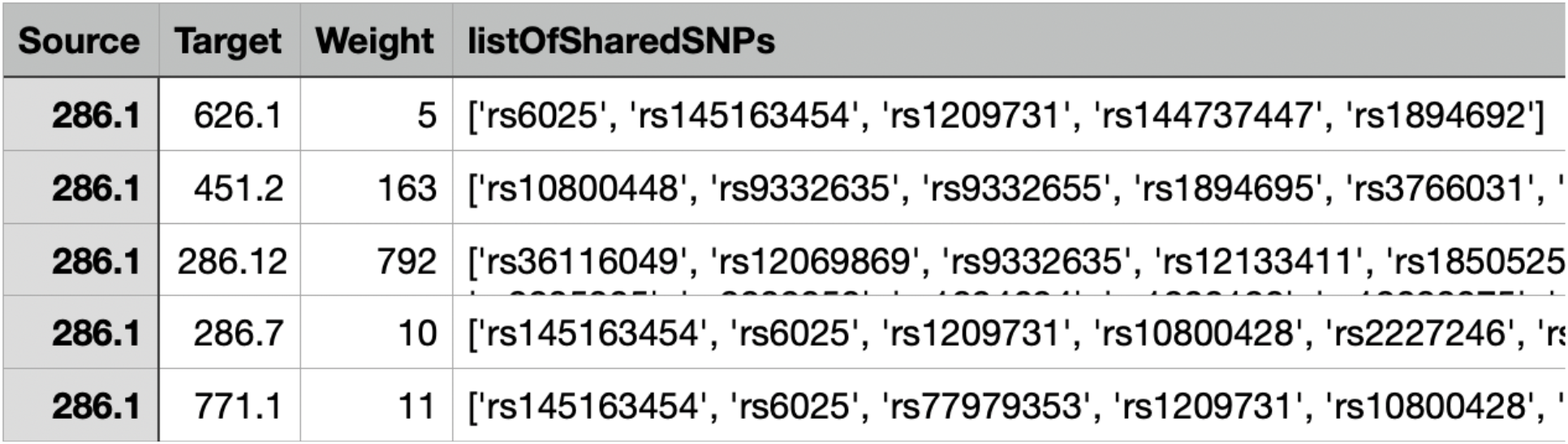
An example edge map

#### b) Node Map

a comma-separated-value file of nodes permitted within our disease-disease network. This dataset should include columns for node names as well as any desired attributes to be displayed in the right side of the visualization.

**Figure 2.**
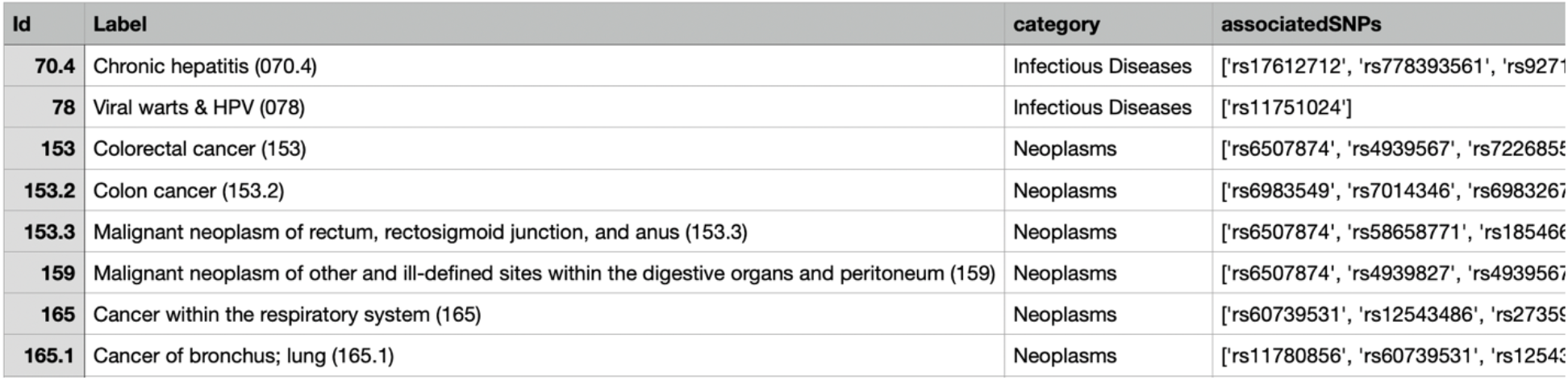
An example node map

#### c) Generating the Node and Edge Maps

The script ***prepDataForGephi.py*** can be used to generate node and edge maps from input summary PheWAS data.

The script requires:

- A directory of summary PheWAS input data (each file should correspond to a different phenotype, and each row of each file should correspond to a SNP found to be associated with the phenotype)
- The name of the column corresponding to SNP name in the input data files
- The delimiter used in the input data files

Note that the input files need to include at least a column corresponding to SNP ID in order for this script to function. No other parameters are essential for the script to run. If the column name for phenotype ID is not provided in input, the script will use the file names themselves as phenotype names.

The script will start off by filtering data if desired: p-value, minor allele frequency, and case count thresholds can all be applied. To include these filters, simply include the column name as well as the desired threshold as parameters of the script. These filtered data are then processed to create a dictionary mapping each phenotype to a list of its associated SNPs. Finally, these SNPs are used to make node maps. If a node map is provided, the associated SNPs (sorted by p-value, if desired), will be appended to it as a last column. If a node map is not provided, a new CSV file will be generated with phenotype and associated SNPs as its two columns. Data are also automatically converted into an adjacency matrix for nodes (the edge map).

Filtration, as well as creation of the node map and the edge map are fully optional. Users can choose to generate these files based upon their inclusion of input and output parameters when calling the python script: 1) if the variable name and threshold value are not specified, data will not be filtered by that variable 2) if an edge map output path is not specified, the edge map will not be generated 3) if a node map output path isn’t specified, the node map will not be generated. For Phecode-based PheWAS data, an input node map that can be used is the “Phecode Definitions 1.2” file: https://phewascatalog.org/phecodes. Note that in this case, phenotype IDs in the input data must match the phenotype ID of the input node map file in order for a node map to be generated.

If all functionality is applied, the output of this script includes two CSV files: the edge map and the node map. Here are a couple examples of how to call “prepDataForGephi.py”:

### Example Usages

1. **Basic (no filtration, input node map, or parameter names beside SNP ID):** python prepDataForGephi.py --input-data “./subset/” --edgemap-output “./subsetEdgeMap.csv” --nodemap-output “./subsetNodeMap.csv” --snp-name “ID” --delim ““
2. **Including all possible parameters:** python prepDataForGephi.py --input-data “./subset/” --edgemapoutput “./subsetEdgeMap.csv” --nodemap-input “./phecodeDefinitions.csv” --nodemap-output “./subsetNodeMap.csv” --maf-threshold “0.01” --casecount-threshold “200” --pvalue-threshold “1e-8”-phenotype-name “phenotypeID” --snp-name “ID” --maf-name “af” --casecount-name “num_cases” --pvalue-name “pval” --delim ““

### d) Creating a Network in Gephi

**Figure 3.**
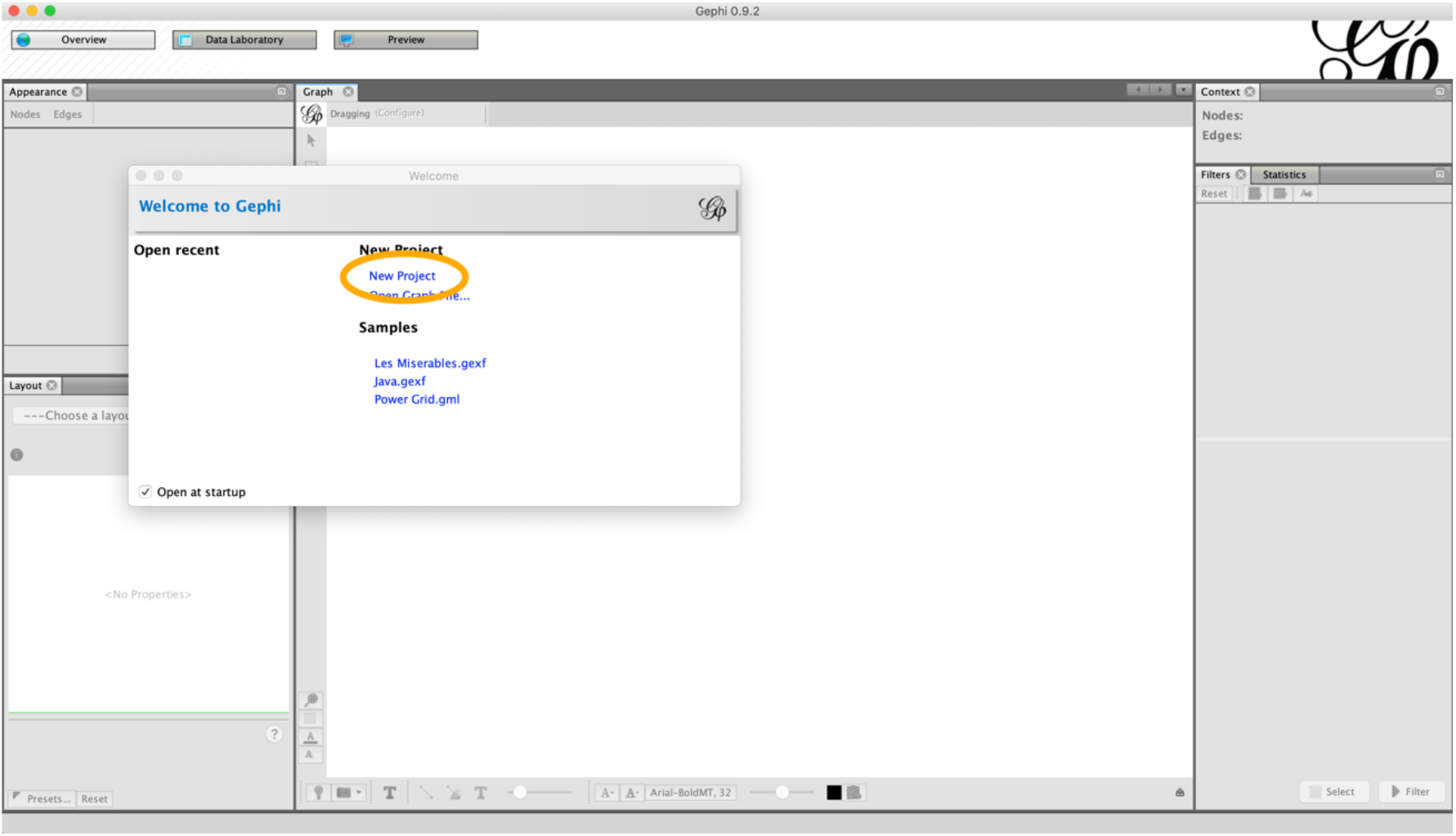
Homepage for Gephi upon initialization. Click on “New Project” to start a new map.

1. Import nodeMap.csv as a Node table. Make sure it is undirected, make sure to add it to the existing workspace, and avoid including extraneous variables

**Figure 4.**
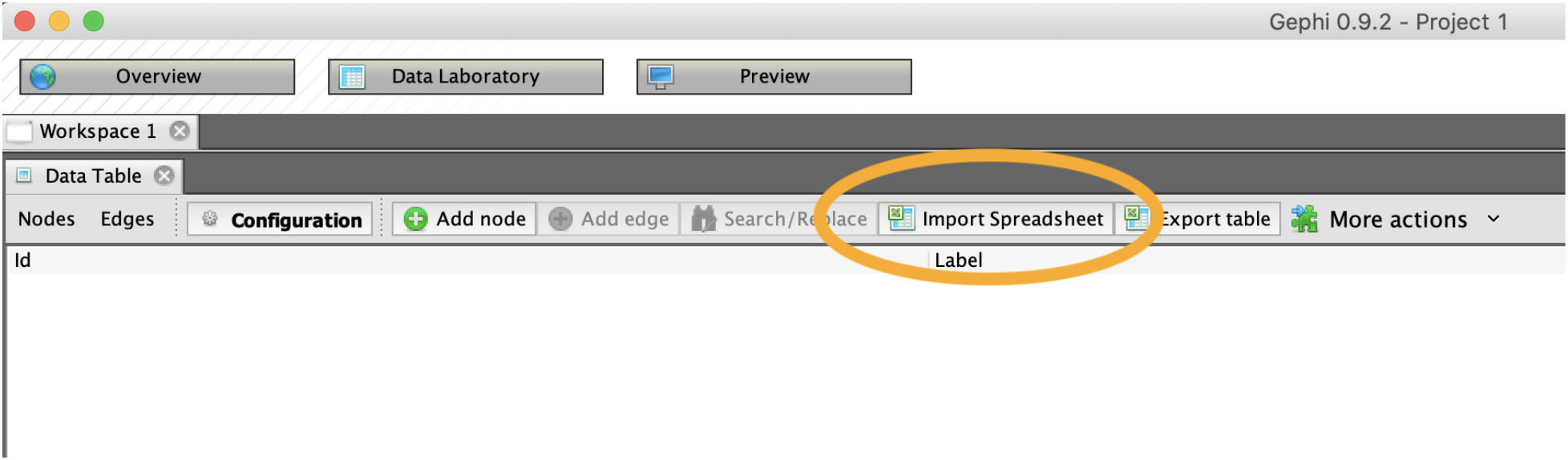
Use the “Import Spreadsheet” button to upload both your node and edge maps.

**Figure 5.**
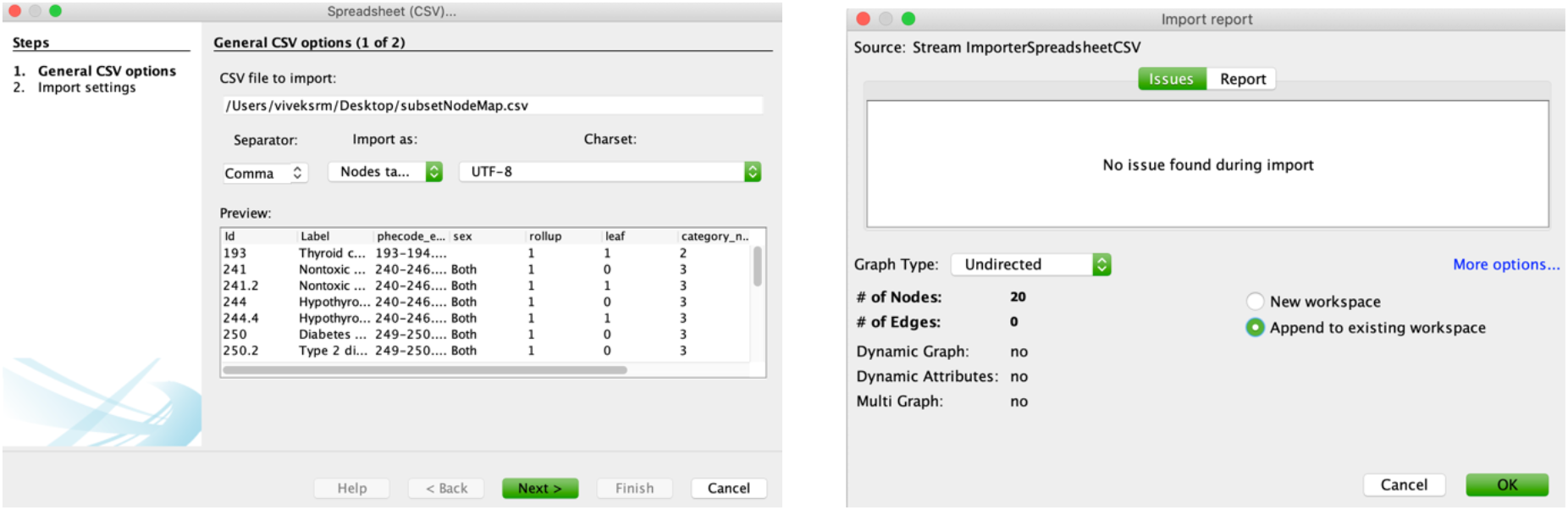
Layout for Node Map upload.

**Figure 6.**
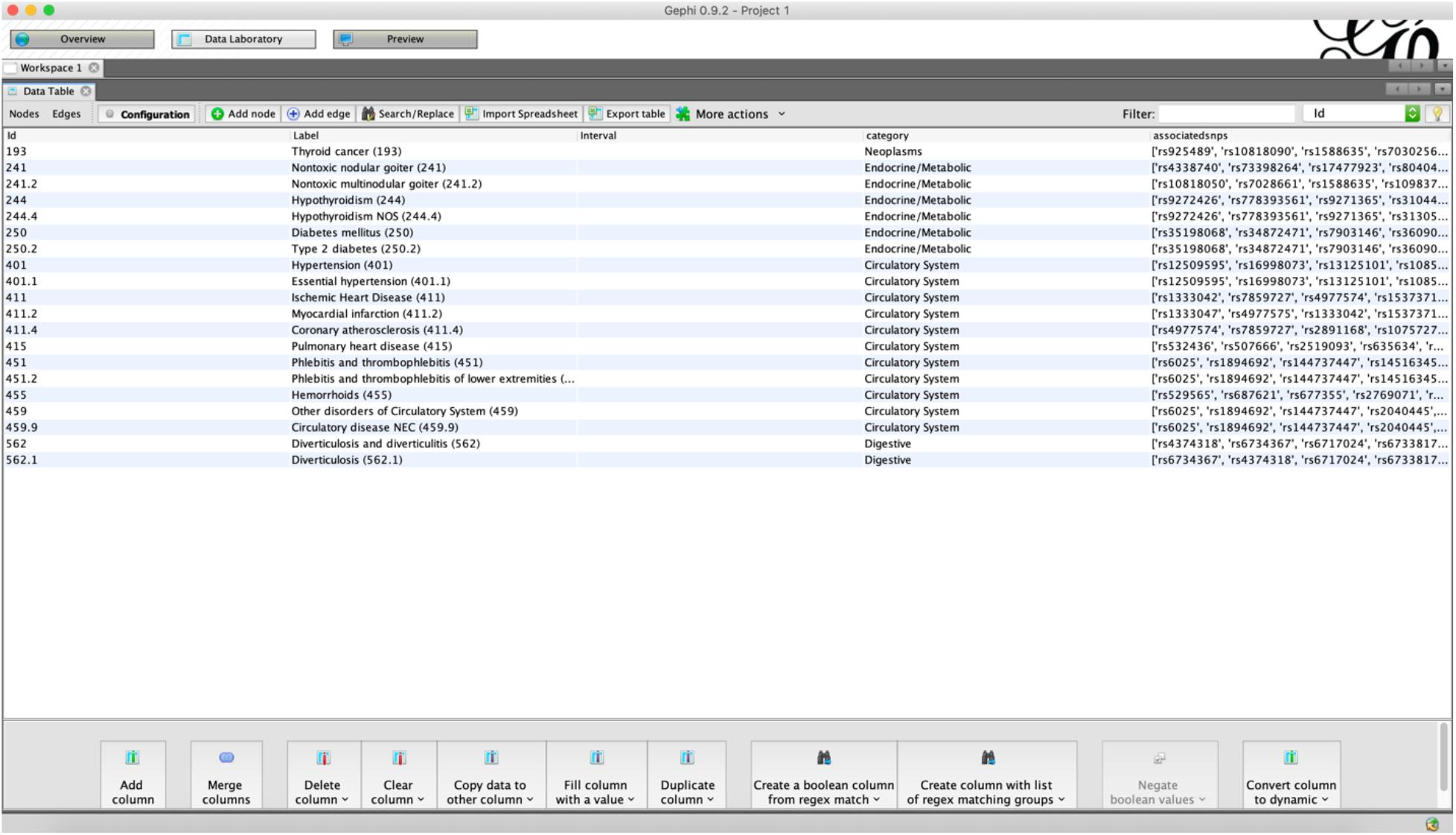
Results of uploading a node map into Gephi

2. Import edgeMap.csv as an Edge table. Make sure to add it to the existing workspace.

**Figure 7.**
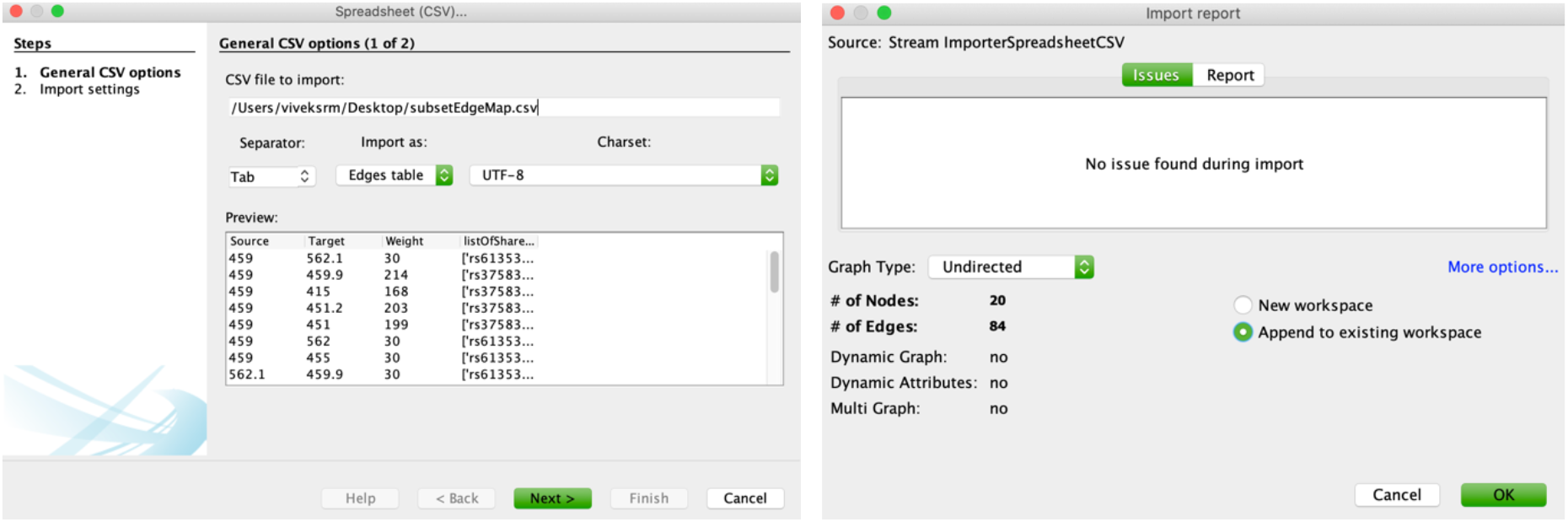
Steps to upload an edge map into Gephi

3. Click on overview to see the current network.

**Figure 8.**
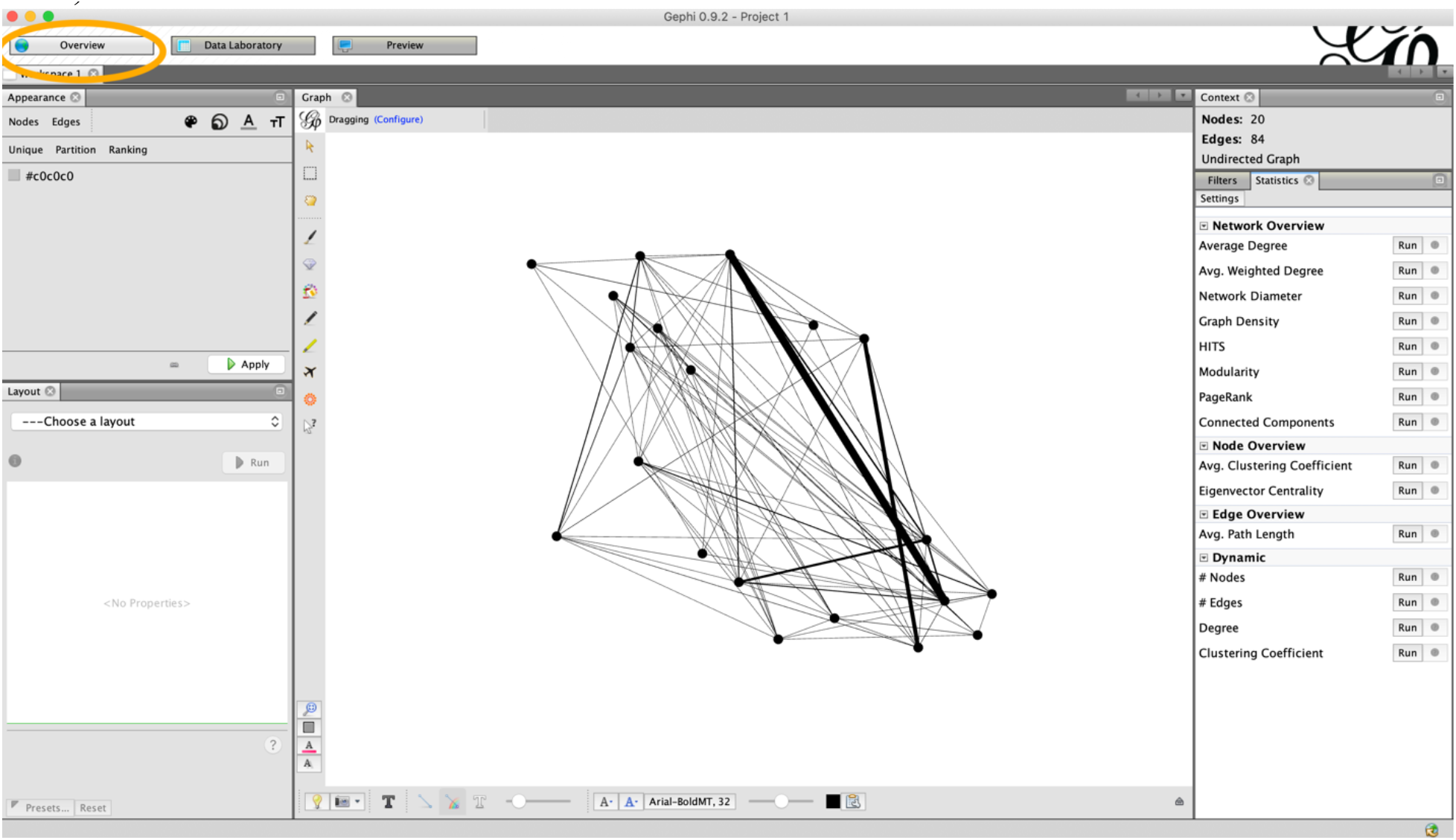
The map can be visualized by clicking “Overview” after uploading the node and edge maps in the Data Laboratory.

4. Using the Statistics menu at the right of the screen, calculate the degrees of all nodes. If you desire, you can navigate back to the Data Laboratory, select nodes that have degree 0, and delete them so that your map looks cleaner.

5. Calculate the rest of the node statistics (and re-calculate degree if you deleted nodes) by clicking “Run” next to each network statistic in the Statistics menu.

**Figure 9.**
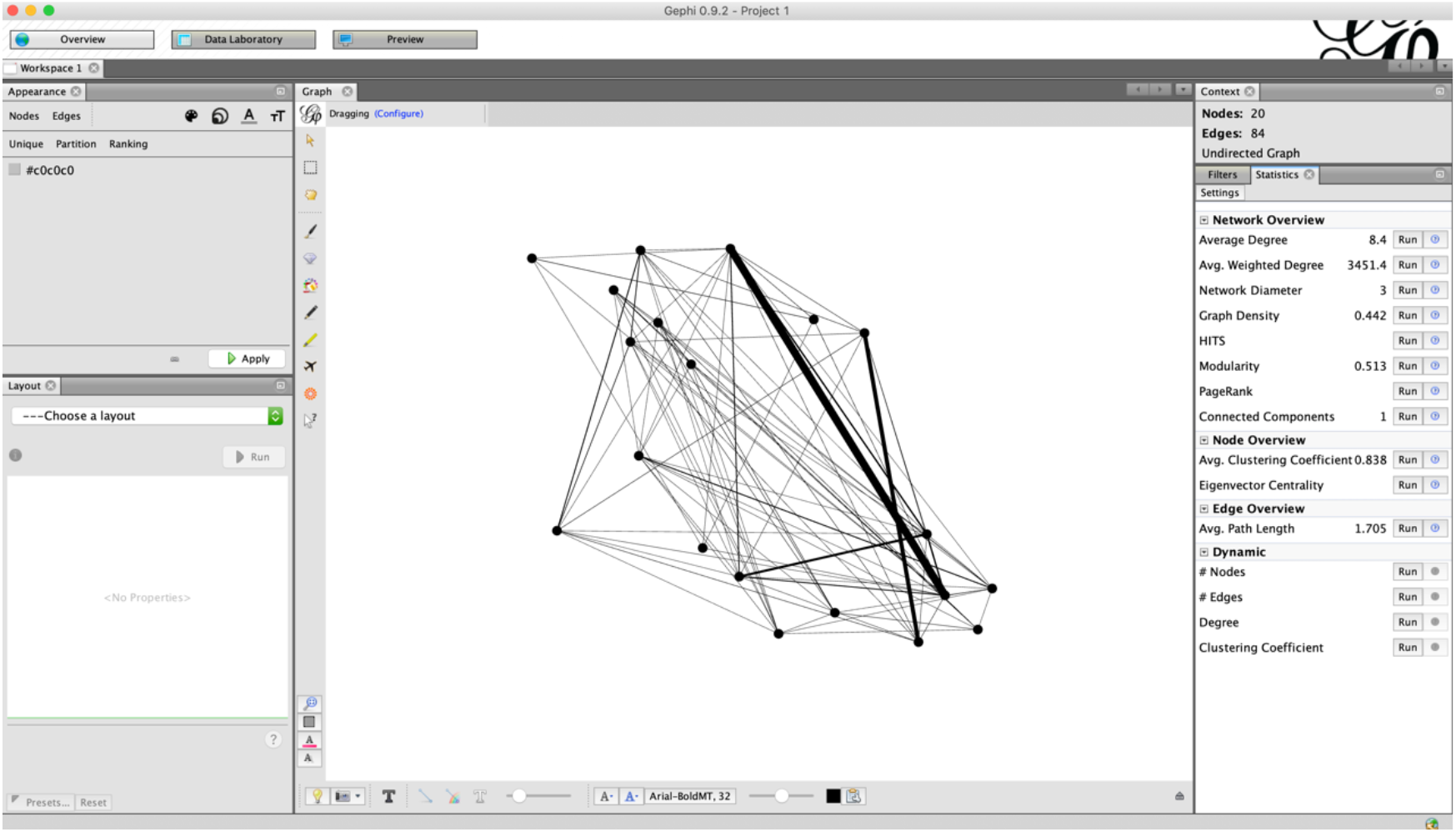
Use the statistics menu on your map by clicking run for each of the options at the right of the screen

6. Fit your desired layout to your network. Points can also be dragged around to yield different shapes.

**Figure 10.**
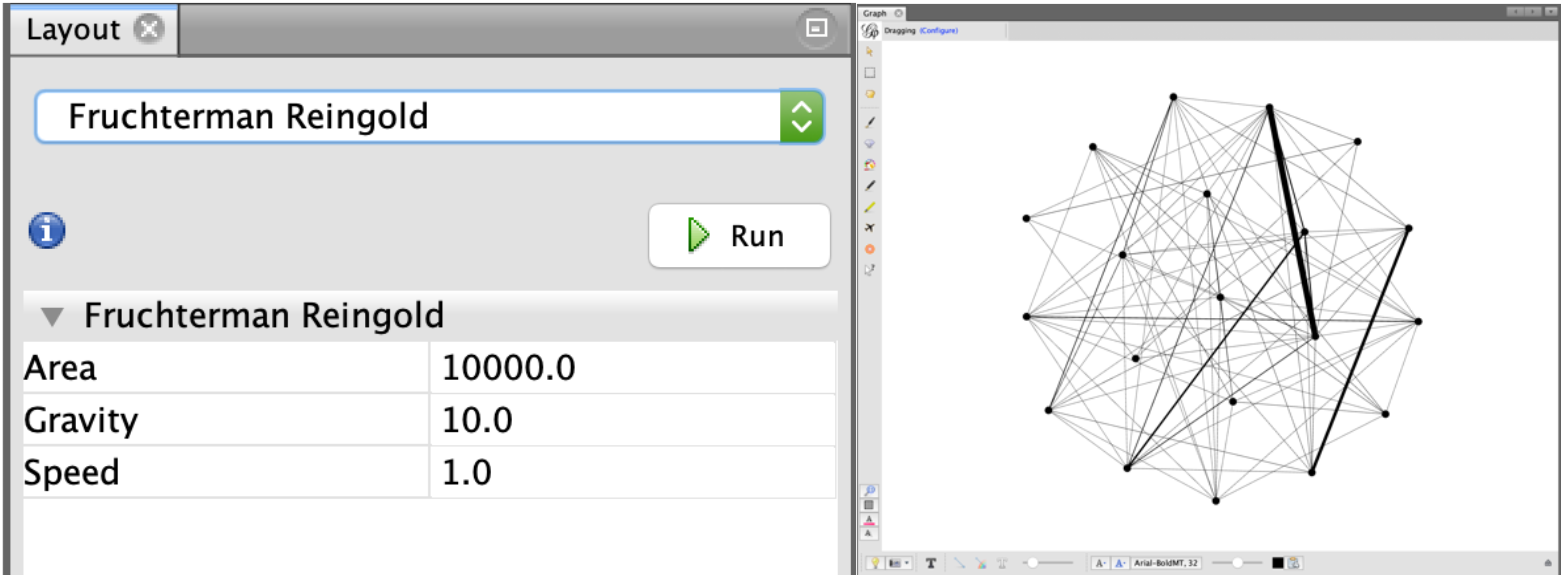
An example of using the Fruchterman Reingold layout to reorganize your network.

7. Color nodes by desired variable (i.e., disease category). Size nodes by desired variable (i.e., node degree)

**Figure 11.**
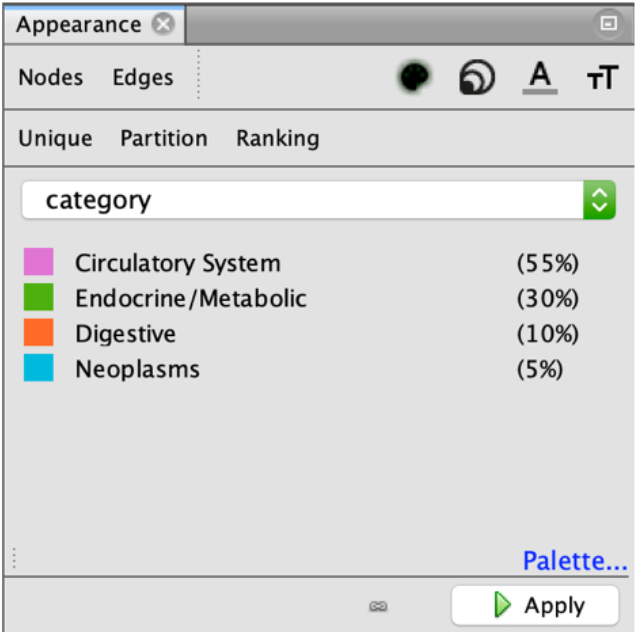
The appearance menu can be used to size and color nodes. To color a node by a variable, click “Node”, click the painter’s palette, click “Ranking,” and then select your desired variable. To size a node, click “node,” click the concentric circles, click “Ranking,” and then select your desired variable.

8. Scale edges up to their full capacity to make sure they are visible after exporting

**Figure 12.**
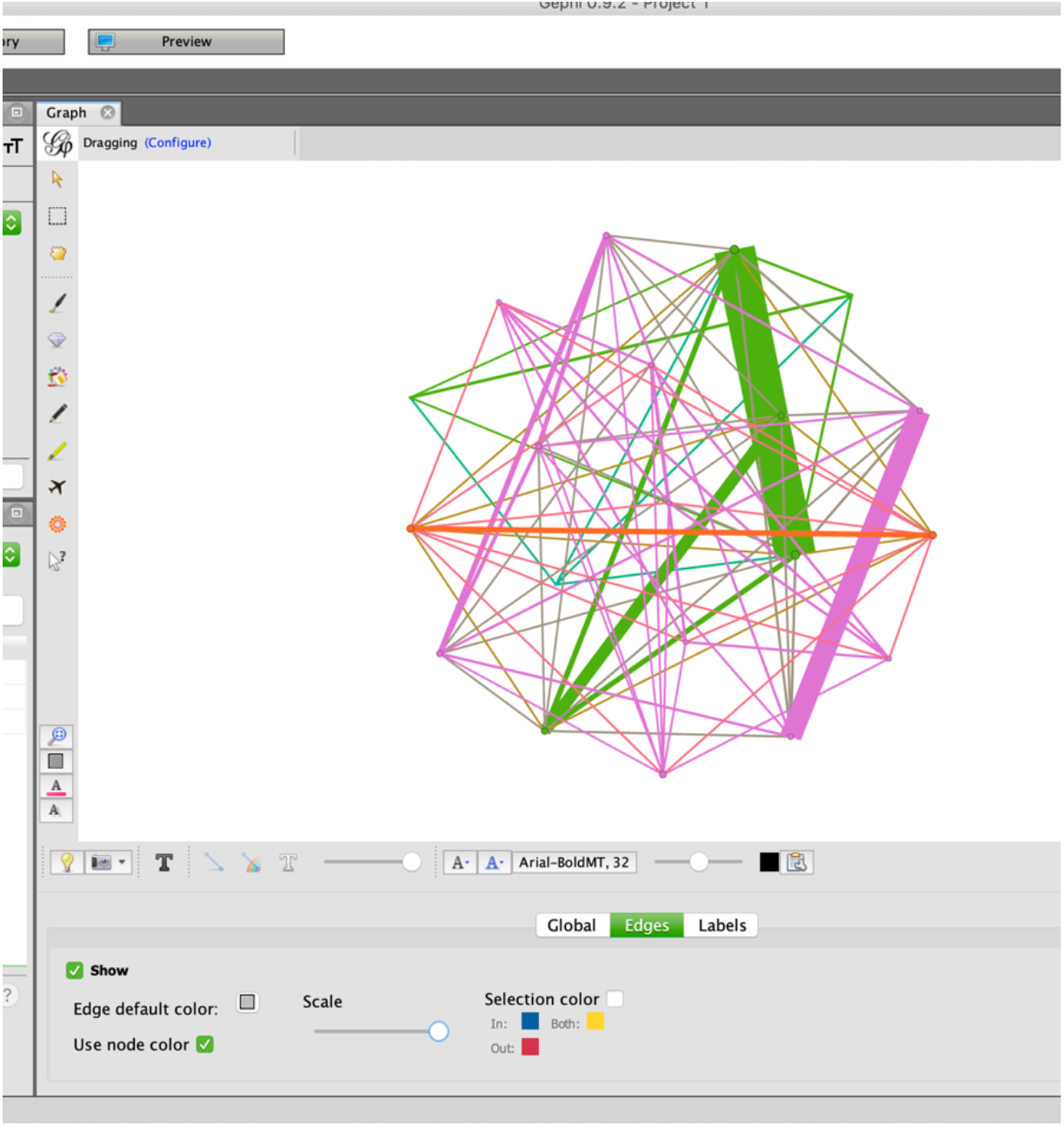
Scale the edges all the way up so that they will remain visible after exporting the map.

9. Export the graph to “Graph file” format. The output of export will be a single “data.json” file. Make sure that the disease category column name is labeled as “Category” (this is case-sensitive). “data.json” can now be plugged into the NETMAGE software for the generation of a corresponding disease-disease network. Data can also be saved in GEXF or XML format from Gephi.

**Figure 13.**
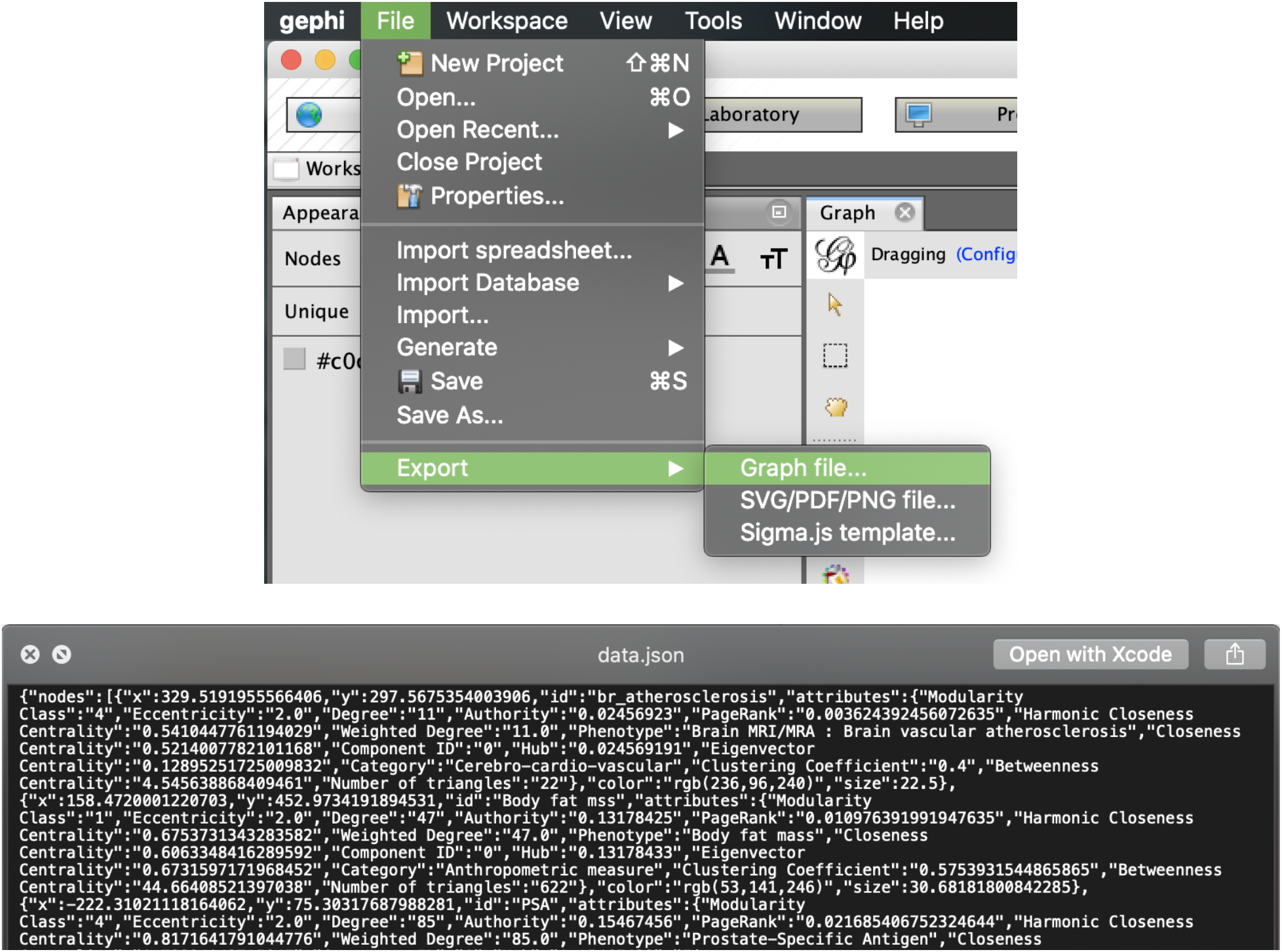
Exporting the map as a “Graph file” will automatically safe the information into a file called “data.json.” This output can be plugged into NETMAGE for the visualization of the network.

### Interacting with the network online

This second section of the guide provides a brief description of how to interact with the resulting disease-disease network produced by NETMAGE.

- Use the drop-down menu to select a variable by which to search the network map
- If the variable selected is continuous, a 2-input slider will appear, allowing users to specify a range of values they want to be permitted. The values allowed in this slider will vary depending on the range of values for the chosen variable in the dataset
- If the variable selected is categorical, a text-box input will appear, allowing users to search for nodes that match the search term

- The Group Selector can be used to display nodes within each category. Nodes are colored as well by the variable chosen for category.
- Any node within the graph can be selected – hovering over a node will show the node’s name
- Selection of a node will display the chosen node and all first-degree neighbors.
- The right side of the visualization will then display a variety of network statistics, such as Modularity Class, Eccentricity, and Degree, as well as other variables included by the user (i.e associated SNPs)

**Figure 14.**
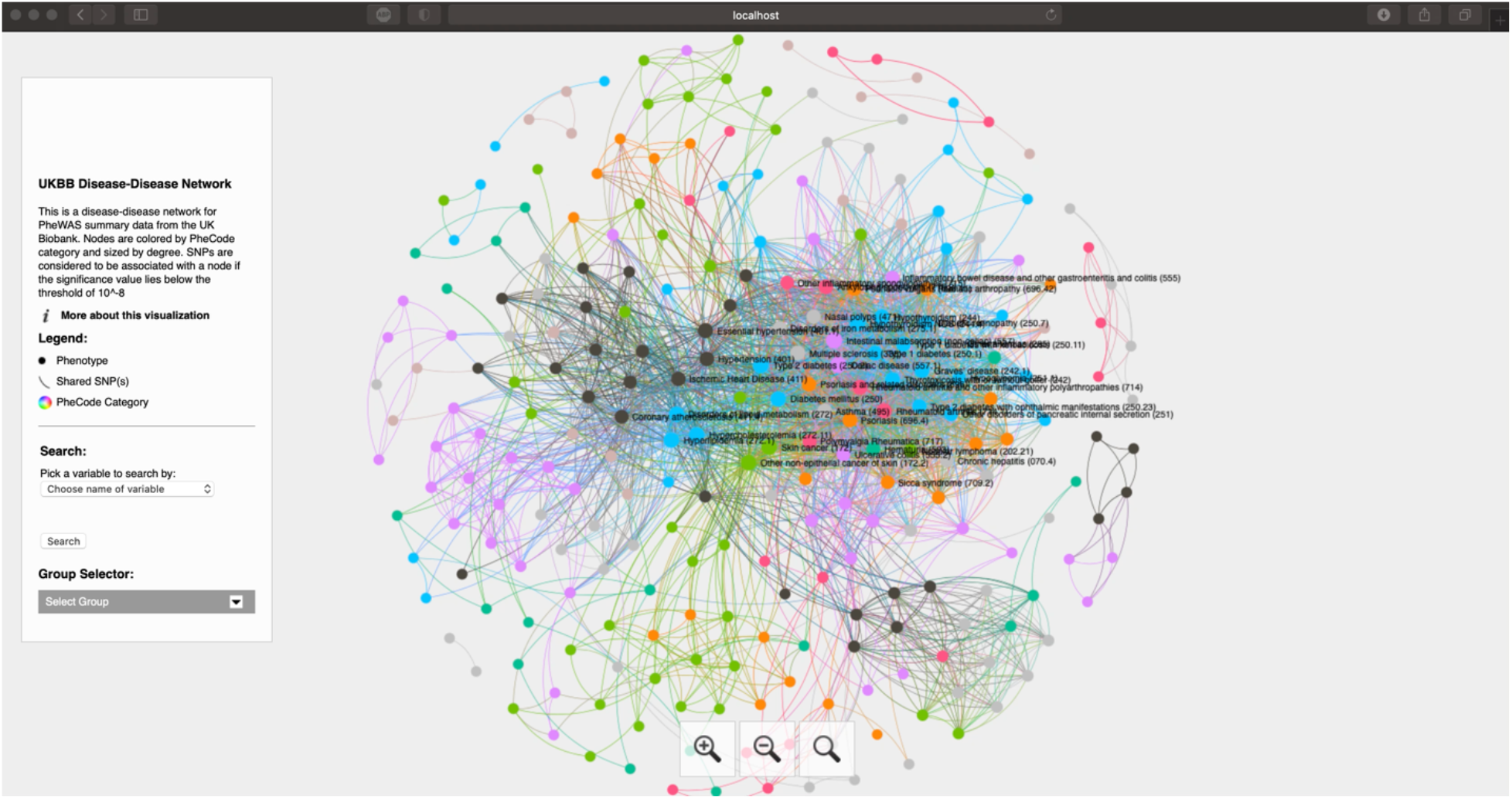
A video demonstration of how to interact with networks produced by NETMAGE.

## Notes

### Competing Interest Statement

The authors have declared no competing interest.

